# Plant stem cell organization and differentiation at single-cell resolution

**DOI:** 10.1101/2020.08.25.267427

**Authors:** James W. Satterlee, Josh Strable, Michael J. Scanlon

## Abstract

Plants maintain populations of pluripotent stem cells in shoot apical meristems (SAMs), which continuously produce new aboveground organs. We used single-cell RNA sequencing to achieve an unbiased characterization of the transcriptional landscape of the maize shoot stem-cell niche and its differentiating cellular descendants. Stem cells housed in the SAM tip are engaged in genome integrity maintenance and exhibit a low rate of cell division, consistent with their contributions to germline and somatic cell fates. Surprisingly, we find no evidence for a canonical stem cell organizing center subtending these cells. In addition, we use trajectory inference to trace the gene expression changes that accompany cell differentiation. These data provide a valuable scaffold on which to better dissect the genetic control of plant shoot morphogenesis.

---

Unlike animals where organogenesis is typically completed in juvenile stages, plants initiate new organs throughout the lifespan via the persistence of pluripotent stem-cell populations long after embryogenesis. These stem cells are housed within the shoot apical meristem (SAM), which gives rise to all of the above ground organs of the plant (*1*). Canonical descriptions of SAM organization in flowering plants include a stem-cell niche within the central zone at the SAM tip, subtended by the stem-cell organizing center, and a peripheral zone surrounding the SAM flank that provides initial cells for organogenesis. The mechanisms that underlie stem-cell niche organization and maintenance, and how cells attain differentiated fates remain fundamental questions in plant development.

Class I *KNOTTED-LIKE HOMEOBOX* (*KNOX*) genes broadly promote indeterminate cell identity in vascular plants (*2*); *KNOX* downregulation is a marker of cell differentiation and comprises an initial step in lateral organ identity acquisition at the SAM periphery. In Arabidopsis, stem-cell homeostasis is achieved via a negative feedback loop wherein the activity of the stem-cell-organizing transcription factor WUSCHEL (WUS) is repressed by binding of the small secreted peptide CLAVATA3 (CLV3) to the CLAVATA1 (CLV1) receptor (*3*). The canonical CLV-WUS signaling pathway and other receptor-ligand signaling complexes regulating WUS-mediated control of shoot meristem size are identified across the flowering plants (*3*).

In order to better understand the spatial organization of the SAM and the process of cell differentiation during plant development, we took a single-cell transcriptomic approach to achieve an unbiased sampling of cell types from the maize (*Zea mays spp. mays*) SAM and seedling shoot apex unimpeded by prior histological assumptions. Improved protocols for the isolation of living plant stem cells enabled this first, to our knowledge, single-cell transcriptomic analysis of a vegetative shoot meristem in any plant. Two zones of cell identity are identified within the maize SAM: (1) a slowly-dividing stem-cell domain at the SAM tip expressing genes with functions in genome integrity, and (2) a subtending population of cells undergoing transit-amplifying divisions. Surprisingly, although the CLV-WUS stem-cell homeostatic pathway is well described in a diverse array of angiosperm SAMs and in the inflorescence and floral meristems of maize (*3*), we do not find evidence for a stem-cell organizing center expressing *WUS* in the maize SAM (*4*). In addition, we use trajectory inference to identify dynamic gene expression patterns correlated with cell differentiation and ultimate cell fate in the seedling shoot. We find that during both early and later stages of seedling shoot ontogeny, a similar set of genes is deployed to mediate the transition from indeterminate to determinate cell fate during plant development.

## Results

### Single-cell transcriptomic approach for the analysis of maize vegetative SAM cells

Single-cell transcriptomic analyses of plant cells require the preparation of protoplasts, viable cells whose rigid, cellulosic cell walls are enzymatically removed. Previously, failures in recovery of viable protoplasts from SAM-enriched plant tissues has presented an obstacle to scRNAseq analyses of shoot meristems (*5*). To achieve a higher rate of viable cell recovery, we supplemented the protoplasting solutions with L-Arginine (L-Arg), which modestly enhanced cell viability (Fig. S1A). This finding was consistent with previous reports of enhanced cell viability of oat (*Avena sativa*) coleoptile protoplasts cultured in media supplemented with L-Arg (*6*). Increasing the pH of the media further enhanced protoplast viability (Fig. S1B), in keeping with prior studies showing that the *in vivo* pH of SAM tissue in the herbaceous plant *Chenopodium rubrum* is two orders of magnitude more alkaline than typical plant protoplasting solutions (*7*). Together, these modifications to our protoplasting protocol improved cell viability between 10-30 fold, depending on the tissue.

To capture cells from the microscopic seedling SAM, we manually harvested protoplasts from dissected apices comprising the SAM plus the two most recently-initiated leaf primordia (SAM + P2). After filtering, six biological replicates captured a total of 327 cells for single-cell RNAseq (scRNA-seq) analyses (Fig. 1A, S2). We first used *k*-means clustering to classify transcriptionally similar cells, adding a number where appropriate to reflect subpopulations within a given cell type. Next, we performed dimensionality reduction using Uniform Manifold Approximation and Projection (UMAP), which plotted the seven resulting clusters corresponding to major cell types derived from the epidermis, primordia, and vasculature, along with indeterminate cell types from the SAM (Fig. 1B, S3). Owing to the abundance of cycling cells in the SAM + P2 tissues (49% in S/G2/M phase), we regressed out variation contributed by the cell cycle on cell clustering (Table S1). Intriguingly, instead of forming discrete, isolated cell clusters, the majority of SAM-enriched cells exhibited a continuum of intermediate identity states (Fig. 1B), suggesting that differentiation is highly dynamic and continuous throughout the maize shoot apex.

**Fig. 1.**
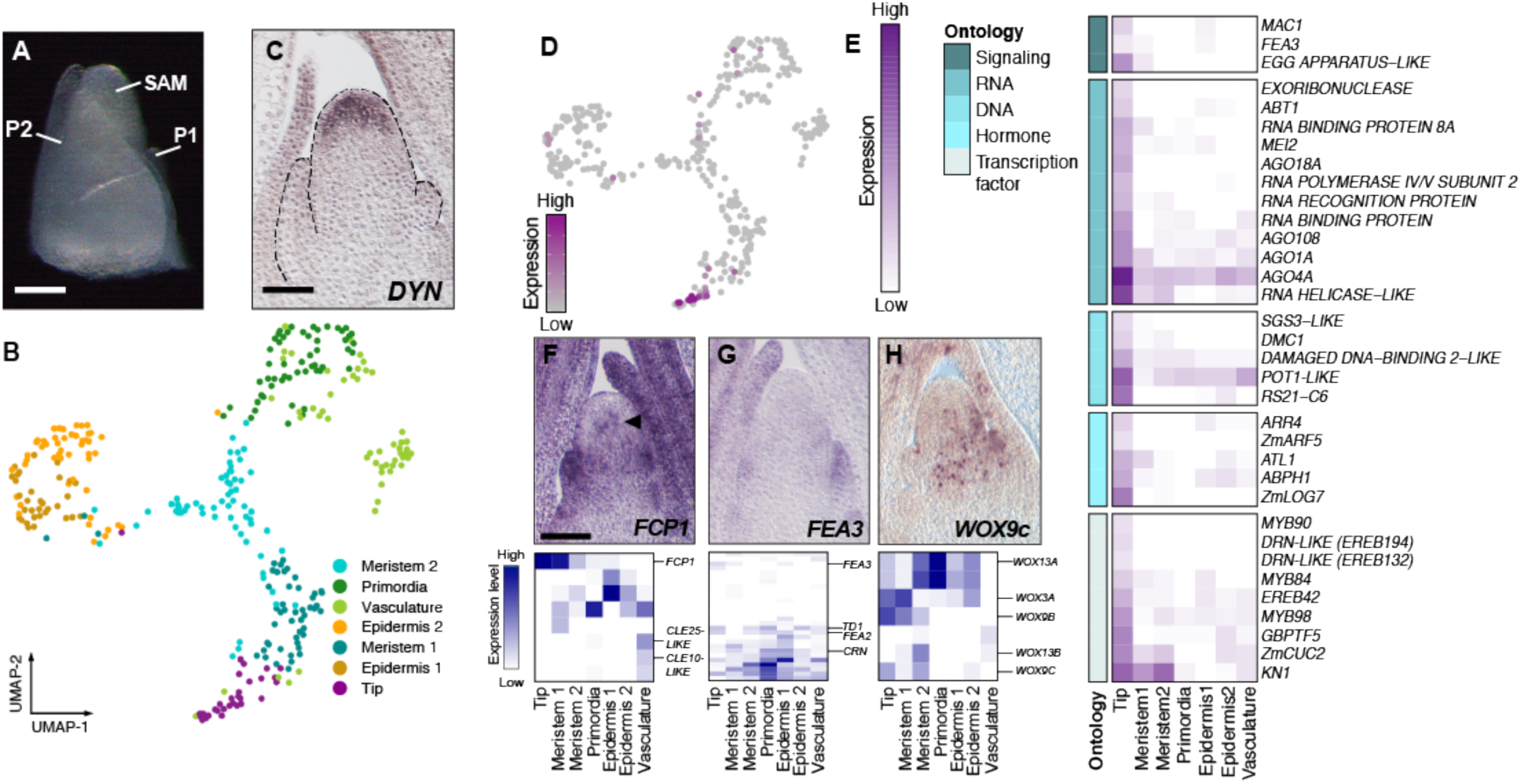
Transcriptomic signatures of stem-cell identity and maintenance in the maize SAM. (A) Cells were isolated from the SAM plus two most recently initiated leaf primordia (SAM + P2). (B) Dimensionality reduction and cell classification for cells in the SAM + P2 dataset. (C) RNA *in situ* hybridization with antisense probe to *DYNAMIN* in medial longitudinal section of the SAM. (D) *DYN*, a marker for the SAM tip and putative stem cell domain, in the SAM + P2 dataset correlates with the putative tip cell population. (E) Heatmap of select differentially expressed genes in the tip domain grouped based on functional ontology. (F-H) RNA *in situ* hybridization with antisense probes to *FCP1* (F), *FEA3* (G) and *WOX9c* (H) (top) and DE analysis along with paralog gene expression in the SAM + P2 dataset (bottom). Scale bars = 100 *µ*m.

### Characterization of the maize stem-cell niche

We first attempted to identify signatures of the stem cell population of the maize SAM. The tip of the maize SAM is thought to house the stem-cells (*4*, *8*), which are essential for the generating the above-ground somatic tissue of the maize plant as well as cells that give rise to the germline. Our cell clustering analysis independently identified a transcriptionally distinct cell population in which *DYNAMIN* (*DYN*), a previously-identified marker of tip cells in the maize SAM, was upregulated (*4*). We therefore designated cells belonging to this cluster as the putative stem cells residing within the SAM tip and used differential expression (DE) analysis to identify 89 genes preferentially expressed within this population (Fig. 1C-D, Table S2) (*4*). Among these were genes with confirmed or predicted roles in intercellular signaling, small RNA biogenesis, DNA maintenance, response to the plant hormones auxin and cytokinin, and transcriptional regulation (Fig. 1E). Closer inspection of the numerous genes (FDR for GO term enrichment = 0.05; Table S3) involved in RNA biogenesis suggested that these cells are engaged in RNA-dependent gene silencing activities. For example, the stem cell-enriched *SUPRESSOR OF GENE SILENCING3-LIKE (SGS3-LIKE)*, *RNA POLYMERASE IV/V SUBUNIT2*, and *ARGONAUTE4a (AGO4a)* all encode members of the RNA-directed DNA methylation pathway that maintains heterochromatin at repetitive, retrotransposon-enriched, maize genomic regions (*9*). Indeed, maintenance of heterochromatin is likewise essential for the genomic stability and homeostasis of stem-cell populations in animals (*10*). However, unlike animals where germline cells are specified and sequestered during early embryonic stages, plants lack a segregated germ cell lineage during vegetative stages of development (*11*). Upregulation of genes involved in DNA repair related processes, such as a *PROTECTION OF TELOMERES1-LIKE (POT1-LIKE)* and a *DNA-DAMAGE BINDING2-LIKE* gene may reflect the advantage of maintaining high genomic fidelity among cells that have the potential for both somatic and germ cell fate (*12*). Collectively, these data suggest that cells in the maize SAM tip are engaged in genome protective functions consistent with their plant stem-cell identity.

### Divergence in SAM stem-cell regulation

We next sought to analyze the cell-specific expression patterns of regulators of stem-cell maintenance within the SAM + P2 tissues. *FON2-LIKE CLE PROTEIN1* (*FCP1*) was the only *CLV3-*like ligand-encoding transcript detected in meristem tip cells (Fig. 1F). RNA *in situ* hybridization identified weak expression of *FCP1* just below the SAM tip, as well as the originally reported expression in the SAM periphery and leaf primordia (*13*). The FCP1 peptide-ligand is perceived by the FEA3 receptor to repress stem-cell identity (*13*). *FEA3* transcripts show low and diffuse accumulation in the SAM periphery, and heightened expression in leaf primordia (Fig. 1G). Other maize transcripts encoding predicted leucine-rich repeat (LRR) receptors exhibited similar accumulation patterns (Table S4), with higher expression in the SAM periphery and primordia but a lack of a strong SAM-specific expression profile as is seen for *CLV1* in Arabidopsis (*14*). This may reflect a role for LRR receptors in inhibiting stem-cell identity outside of the SAM tip domain, as described previously for the FCP1-FEA3 ligand-receptor system (*13*). Notably, mutations in the *CLV1* and *CLV2* homologs of maize cause enlarged inflorescence meristems, but do not affect vegetative SAM size (*3*).

In Arabidopsis, the stem-cell promoting transcription factor WUS is negatively regulated by CLV1-CLV3 function, to control the size of the meristematic stem-cell pool (*3*). WUS is mobile and is expressed in the organizing center below the stem-cell domain from where it is trafficked to promote stem-cell fate in the SAM tip. A similar, stem-cell organizing *Zm*WUS function is described in the maize inflorescence meristem (*3*). However, no *ZmWUS*-expressing cells were identified in the SAM, although transcripts of several maize *WUS* homologs including *ZmWOX3a*, *ZmWOX9b* and *ZmWOX9c* were detected in the SAM (Fig. 1H). The single Arabidopsis *WOX9* homolog promotes cell proliferation in meristematic tissues upstream of *WUS* function, but belongs to a more ancient, functionally-divergent WOX clade that lacks the repressive WUS-box (*15*, *16*). Overall, this suggests that the maize co-orthologs *WOX9b* and *WOX9c* are unlikely to be functionally homologous to *WUS*. Moreover, although *WOX3A* does encode a WUS-box and is detected in the SAM (Fig. 1H), its expression pattern is not cell-type specific. Thus, our data identify no candidate *WOX* gene expressed in the maize SAM that is likely to function as a stem-cell organizing center, homologous to *WUS* in Arabidopsis and *ZmWUS* in the maize inflorescence meristem. Together, these results suggest that the canonical CLV1-CLV3-WUS pathway has been bypassed in the maize SAM, with changes in the spatial expression patterns of corresponding maize paralogs as a defining feature.

### Characterization of a SAM core region

To further investigate the organization of the maize SAM we examined gene expression patterns among cells derived from a recently-reported domain situated in the center, or ‘core’ region, of the SAM, which is marked by the expression of *GRMZM2G049151*, a gene of unknown function (*4*). We identified cells within the core region by transcript accumulation of *GRMZM2G049151*, and used differential expression (DE) analysis to characterize their expression profiles (Fig. 2A-B, Table S5) (*4*). To reiterate our results above, no maize *WOX* genes are DE in the SAM core. In addition, SAM core cells show upregulated expression of the *PLASTOCHRON1* (*PLA1*) gene. *PLA1* promotes cell division and growth in an auxin-dependent manner, and is also expressed within multiple maize organs and tissues (*17*). The auxin-promoted dormancy-associated gene *DRM1* (*18*) and the maize *HAIRY MERISTEM3* (*HAM3*) homolog *GRAS33* were also upregulated in the SAM core, as confirmed by RNA *in situ* hybridizations (Fig. 2C-D, S4B-C). Arabidopsis *HAM* genes are expressed in the organizing center, SAM periphery, and in leaf primordia. *AtHAM* genes promote SAM maintenance through their physical interaction with WUS, and also activate the formation and maintenance of axillary meristems (AMs) that give rise to lateral branches (*19*, *20*). Indeed, both *DRM1* and *GRAS33* show expanded expression in maize AMs (Fig. 2E-F). To determine If *GRAS33* activity in the core domain has a conserved role in maintenance of the maize SAM, we generated *gras32 gras33* double mutant seedlings and analyzed SAM size (Fig. S4A). Compared to the wild-type siblings, *gras32/+ gras33* seedlings and *gras32 gras33* double mutants displayed shorter SAMs (Fig. 2G-I). Notably, only *gras32 gras33* SAMs possessed shorter AMs, likely owing to genetic redundancy between these factors in these branch meristems (Fig. 2J). These data suggest that maize *GRAS32* and *GRAS33* may, like their Arabidopsis homologs, have roles in regulating SAM homeostasis from a stem cell-surrounding region that overlaps the maize SAM core domain. These results further suggest that unlike in Arabidopsis, the maize SAM tip is not subtended by a *WOX*-expressing, stem-cell organizing center. Rather, this SAM core region may be engaged in auxin response and functionally akin to the tip-subtending SAM regions expressing *HAM* genes in Arabidopsis, with maize *HAM* genes having a WUS-independent SAM regulatory function.

**Fig. 2.**
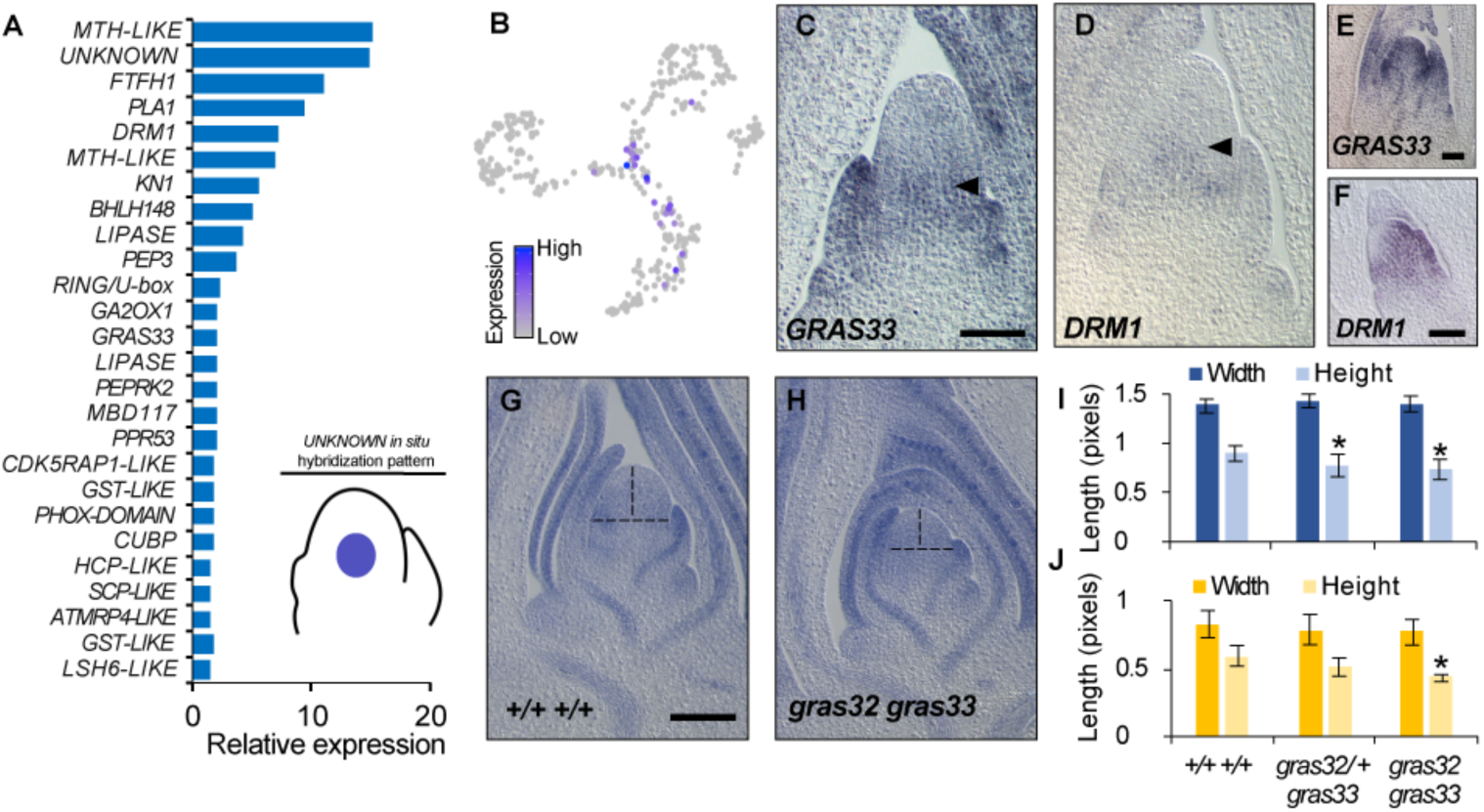
Characterization of the SAM core. (A) Expression of select marker genes in cells positive for expression of the *UNKNOWN* core marker gene. The illustration reflects the published *UNKNOWN* gene expression pattern (*4*). (B) Cells expressing the *UNKNOWN* marker gene in the SAM + P2 dataset. (C-F) RNA *in situ* hybridization of antisense probes to *GRAS32* (C,E) and *DRM1* (D,F) shows transcript accumulation patterns in SAM (C,D) and AM (E,F) medial longitudinal sections. (G,H) Toluidine Blue-O stained medial longitudinal sections of the SAM from normal siblings and *gras32 gras33* double mutants. Vertical and horizontal dashed lines indicate SAM height and width, respectively. (I, J) Quantification of SAM (*n* = 5-8) (I) and AM (*n* = 3) (J) height and width in normal siblings and *gras32 gras33* double mutants (two-tailed Student’s t-test, * *p* < 0.05). Scale bars = 100 *µ*m.

### Rates of cell division are kept low in the SAM stem-cell niche

We next asked whether different maize SAM domains exhibit equivalent cell division rates. Estimations of cell-cycle stage generated during cell-cycle regression (see Methods) indicate that the fraction of SAM + P2 cells in G1 phase decreases as differentiation progresses, indicative of higher rates of cell division (Fig. 3A). For example, the SAM tip population contains a larger fraction of cells in G1-phase than the remainder of the meristem, leaf primordia, or vasculature cell populations, suggesting a lower cell division rate among the stem cells. In order to test this hypothesis, we performed RNA *in situ* hybridization on *HISTONE H3* and *CYCLIN1* transcripts that accumulate in cells at S-phase and G2/M-phase, respectively. We next subdivided medial SAM sections into five proximodistal bins of equal height along the proximodistal axis, and inferred cell division levels by image thresholding on *HISTONE H3* and *CYCLIN1* staining (Fig. 3B-E). Cells in bin 1 that comprises the tip-most region of the SAM consistently had the lowest number of dividing cells. When considered together with the previously described accumulation of transcripts promoting genomic and epigenomic stability in the SAM tip (Fig. 1E), a low cell division rate among the stem-cell population may explain how plants avoid unfavorable increases in genetic load over successive generations in the absence of a segregated germline early in ontogeny. A reduced rate of cell division at the SAM tip is likewise consistent with the low number of cell divisions that are predicted to occur between formation of the maize zygote and the gametophytes contributing to the next generation (*21*). Cells in the remainder of the SAM, beyond the tip, showed higher rates of cell division, similar to transit amplifying cell divisions found in animal stem cell niches (*22*). These proliferative cell divisions in transit-amplifying cells generate the anlagen for determinate lateral organs, obviating the requirement for high levels of stem-cell divisions. Finally, we observed that in AMs, the highest concentration of dividing cells typically occurs closer to the AM tip as compared to the SAM (Fig. 3D). This may reflect changes in the size of the meristem core domain in the AM, in line with our observations of expanded core marker gene expression in AMs relative to the SAM.

**Fig. 3.**
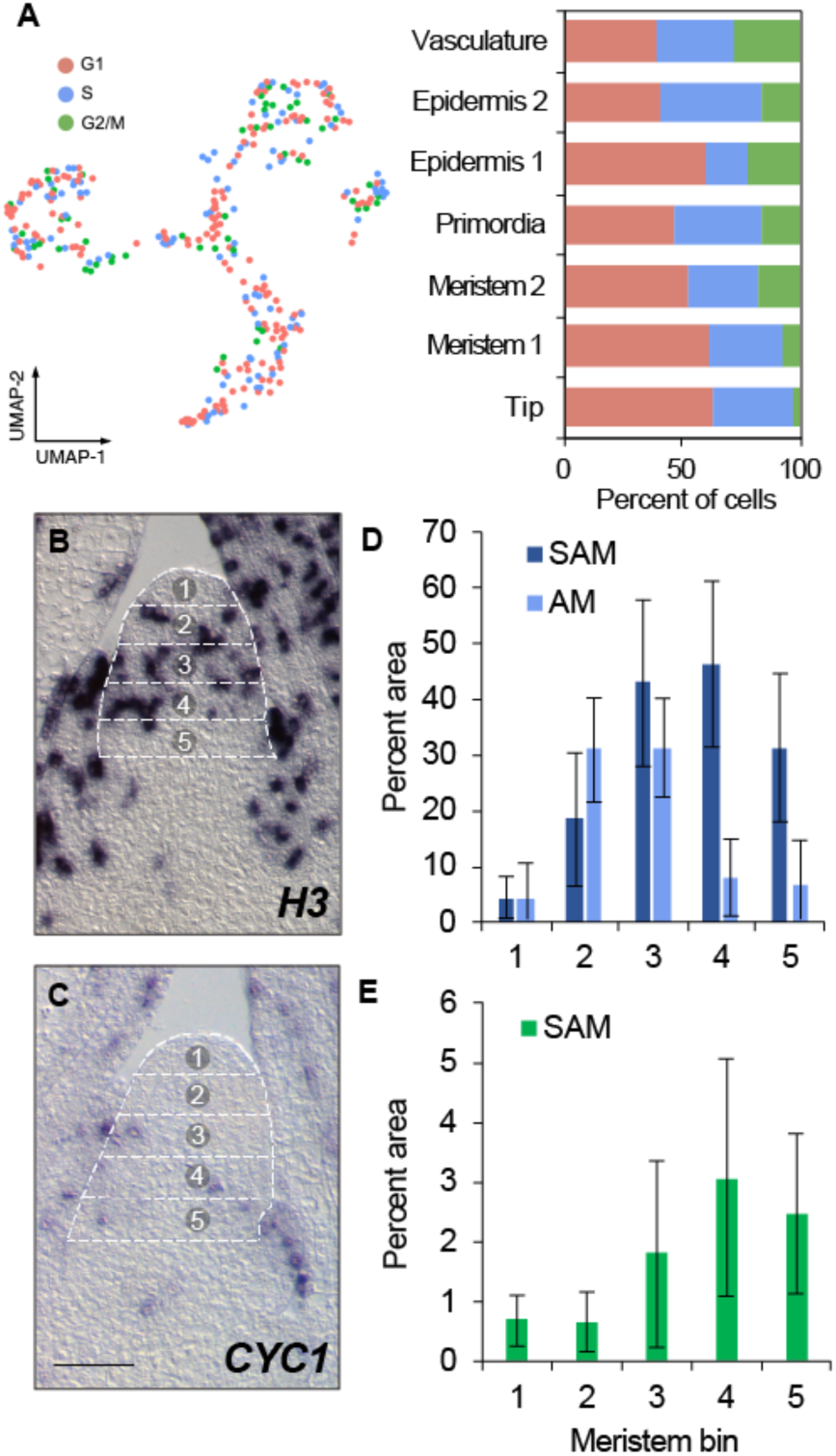
Cell division dynamics throughout the maize SAM. (A) Estimated cell division stage of cells in the SAM + P2 datasets. Bar charts show fraction of cells in each stage among cell clusters. (B,C) RNA *in situ* hybridization with antisense probe to *H3* (B; SAM, *n* = 9; AM, *n* = 4) and *CYC1* (C; SAM, *n* = 7) in medial longitudinal sections of the SAM showing bins (outlined in a dotted line grid and numbered) used for quantification of the proportion of cells in (D) S-phase and (E) G2/M-phase. Scale bars = 100 *µ*m.

### Cell differentiation follows a continuum of transcriptional states

Given that the individual transcriptomes of cells within the maize shoot apex fall overwhelmingly along a continuum of cell differentiation states (Fig. 1B), we aimed to determine the dynamic changes in gene expression that accompany this developmental progression. We applied a principal graph algorithm to identify a branching path among the embedded cellular coordinates in the UMAP projection, which we used to infer the differentiation trajectory of cells in the SAM + P2 dataset. Each cell was then assigned a pseudotime value based on its distance along the resulting path, relative to a specified pseudotime start position within the SAM tip cell population (Fig. 4A). As expected, we found that pseudotime progression is associated with the transition of cells from indeterminate to determinate cell fates; *KN1* and other markers of indeterminate meristematic identity are highly expressed early in pseudotime, whereas genes correlated with lateral organ development such as *GA2OX1* and the YABBY-family transcription factor-encoding gene *DROOPING LEAF1* (*DRL1*) (*23*) show increased expression levels as pseudotime progresses (Fig. 4B). To survey the transcriptional changes associated with cell differentiation over time, we performed differential gene expression analysis to identify transcript accumulation patterns that significantly correlate with pseudotime progression. In total, over 2000 genes exhibited significant changes in expression over pseudotime (Table S6). Hierarchical clustering grouped each transcript according to expression pattern and identified several patterns of transcript accumulation that correspond to particular stages of cell differentiation (Fig. 4C).

**Fig. 4.**
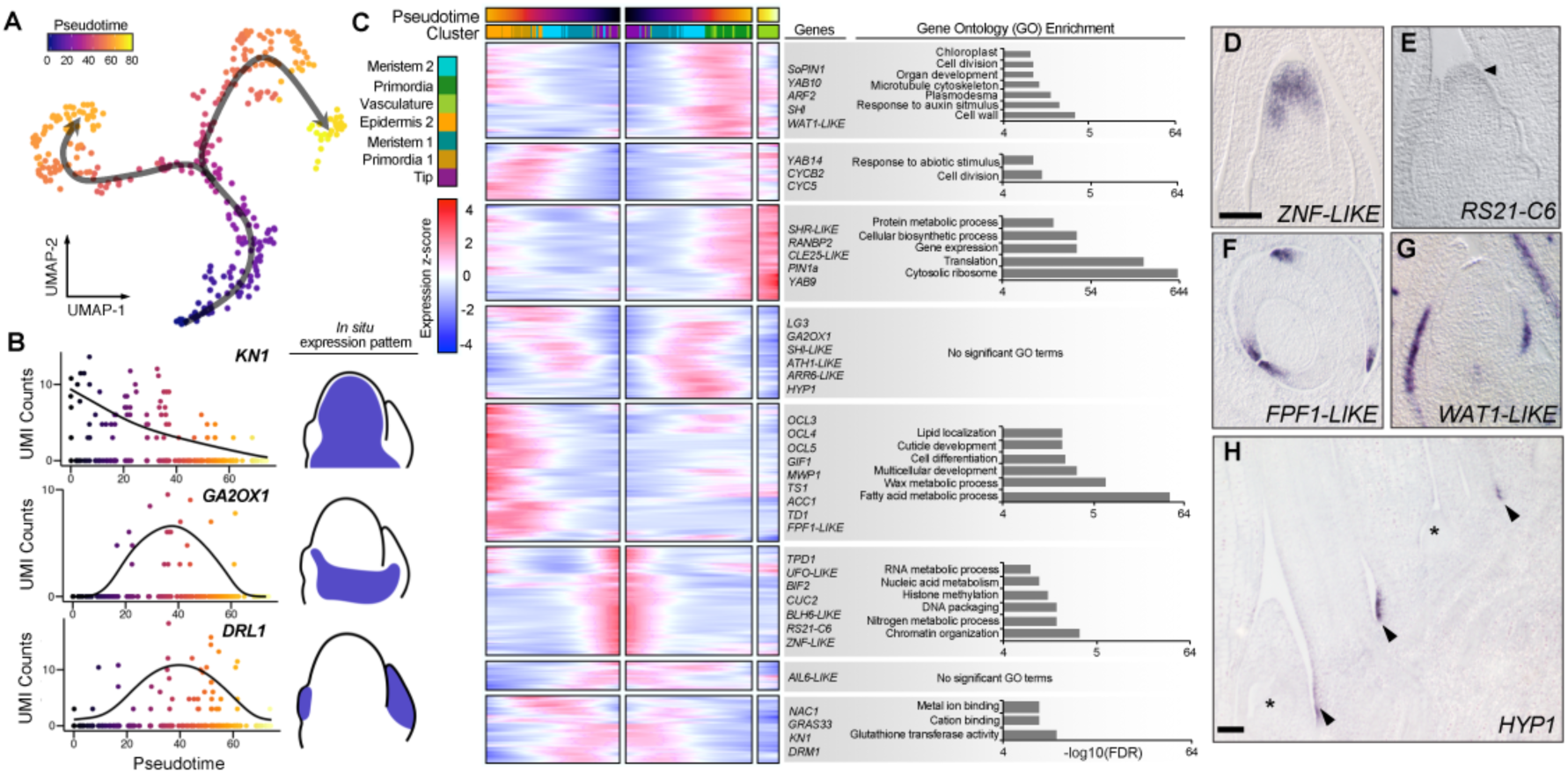
Tracing the gene expression patterns associated with cell differentiation. (A) Pseudotime values and trajectories for cells in the SAM + P2 dataset. (B) Gene expression patterns of published marker genes of cell differentiation over pseudotime and illustrations of their associated transcript accumulation patterns from RNA *in situ* hybridization studies. (C) Heatmap of approximately 2000 genes that show correlated changes in gene expression along the inferred trajectory clustered based on their expression patterns. Cells are mirrored along the central axis prior to the trajectory branch point. Representative genes and significant GO term enrichments for each cluster are shown. (D-H) Transcript accumulation patterns for trajectory-correlated genes showing high expression levels at early, intermediate and late points in the trajectory. RNA *in situ* hybridization of antisense probes to *ZNF-LIKE* and RS21-C6 in medial longitudinal sections of the SAM show early trajectory expression (D,E), *FPF1-LIKE* in transverse section above the SAM and *WAT1-LIKE* in medial longitudinal section of the SAM show late trajectory expression (F,G), and *HYP1* in longitudinal section below the SAM shows intermediate trajectory expression (H). Arrowheads in (E,H) and asterisks in (H) represent transcript accumulation and AMs, respectively.

Early in pseudotime, genes enriched for stem-cell functions in RNA metabolism and chromatin organization give way to genes enriched for glutathione transferase and cation-binding activities as well as expression of *DRM1* and *GRAS33*, which are expressed among transit-amplifying cells. Next, cells progress through a putative boundary domain identity, marked by upregulation of an *ARABIDOPSIS THALIANA HOMEOBOX GENE1-LIKE* (*ATH1-LIKE*) gene, *LIGULELESS3* (*LG3*), and *GA2OX1*. *ATH1* promotes organ boundary formation in *Arabidopsis* and antagonizes activity of the growth phytohormone gibberellic acid, while *GA2OX1* catabolizes gibberellic acid (*24*, *25*). In addition, *LG3* is expressed in specific boundaries during leaf development (*26*). After progressing to this putative shoot boundary domain, the cellular transcriptomes of SAM + P2 cells resolve into either epidermal, or ground and vascular cell identities. The lack of a transcriptionally distinct lineage of undifferentiated epidermal cells (protoderm) early in the trajectory is notable, given that the cell lineage of the outer protodermal layer is separate from that of underlying cells, even within the SAM tip (*1*). Together, this suggests that despite their cell lineage differences, the distinctive transcriptional profiles among cell types in the SAM tip become detectable only after exiting the stem cell niche.

As expected, we found that epidermal cell differentiation correlates with upregulation of the OUTER CELL LAYER homeodomain leucine zipper IV transcription factor-encoding genes that promote epidermal cell identity (*27*). On the other hand, cells fated to become leaf primordia and/or vasculature exhibit upregulation of auxin response genes and transcripts associated with cell wall, chloroplast, and organ development. In addition, primordia and vascular cells are significantly enriched for transcripts related to translation, suggesting a large burst in protein synthesis accompanies leaf initiation and expansion. Selected genes significantly associated with pseudotime progression were examined by RNA *in situ* hybridization to validate their expression patterns along the developmental trajectory (Fig. 4D-H). Two genes expressed at the start of the pseudotime trajectory, *ZINC FINGER DOMAIN-LIKE* (*ZNF-LIKE*) and *RS21-C6*, showed expression in the SAM tip (Fig. 4D,E). Meanwhile, *FLOWERING PROMOTING FACTOR1-LIKE* (*FPF1-LIKE*) and *WALLS ARE THIN1-LIKE* (*WAT1-LIKE*) transcripts are upregulated later in pseudotime and accumulate in the differentiating cells of leaf primordia (Fig. 4F,G). Lastly, *HYBRID PROLINE-RICH PROTEIN1* (*HYP1*) transcripts are upregulated in the pseudotime boundary cluster, albeit detected at later stages of ontogeny (Fig. 4H). Together, these results support the notion that the transcriptomes of differentiating cells are highly dynamic across a continuum defined by pseudotime progression.

### Redeployment of genes patterning determinate and indeterminate cells across ontogeny

After leaf initiation at the SAM periphery, cells continue to proliferate in the leaf proximal region until beyond the P6 stage (i.e. the sixth leaf from the SAM), necessitating continued patterning of indeterminate and determinate zones at the junction of the leaf and stem across ontogeny (*28*). We hypothesized that the transcriptomic signatures of this developmental process would be similar at both early and late stages of leaf development, reflecting the iterative and modular patterning of the plant shoot system. To test this, we isolated cells derived from tissues comprising approximately 3 mm of the maize shoot apex, dissected to include the 6 most recently-initiated leaf primordia plus the SAM (SAM + P6; Fig. S5A). In total, we captured the transcriptomes of over 10,000 protoplasts (in two biological replicates) using microfluidic droplet capture. To identify broad classes of cells, we performed dimensionality reduction and found clusters corresponding to epidermal, vascular, leaf primordial, indeterminate, and cell cycle states (Fig. S5B, S6, S7, Tables S7-9). Cells within our SAM + P6 dataset are overwhelmingly derived from later stages of shoot ontogeny (i.e. P4 to P6), owing to the markedly increased size of these older leaf primordia and associated stems. For example, although we identified indeterminate cells based on their expression of the transcription factor gene *KN1*, its duplicate paralogous genes *ROUGH SHEATH1* (*RS1*) and *GNARLEY1* (*GN1*), as well as the KN1-direct gene target (*GIBERELLIC ACID 2-OXIDASE1*) *GA2OX1* (Fig. S7C) (*25*, *29*) we did not identify cells with transcriptomic signatures of the seedling SAM stem cell niche in our SAM + P6 dataset (Fig. 1E).

We therefore selected a subset of cells spanning this indeterminate-determinate junction in the SAM + P6 dataset to represent cell differentiation at later stages of seedling leaf ontogeny (Fig. S9A). Among these, indeterminate cells expressing *KN1* and a *BLH14* (*BELL1-LIKE HOMEOBOX14*) gene give way to cells occupying a boundary domain, characterized by expression of the maize homolog of *CUP-SHAPED COTYLEDONS2* (*ZmCUC2*) (*26*, *30*) (Fig. S9B). Intriguingly, boundary-region cells were also positive for expression of *WOX9c*, an observation supported by RNA *in situ* hybridization, which suggests a potential non-meristem related function for *WOX9c* (Fig. S9C). Determinate cells were characterized by upregulated expression of the *YABBY*-family transcription factor genes *YAB15* and *DRL1* (Fig. S9C). We again used trajectory inference to assign cells a pseudotime value reflective of their position in the indeterminate-determinate transition and utilized DE analysis to identify genes with pseudotime-correlated expression patterns (Fig. 5A). Overall, we identified approximately 3000 genes that met a stringent significance cutoff (adj. *p* < 1E-100) and compared them to pseudotime-correlated genes in the SAM + P2 dataset. We found that approximately one third of the transcripts showed a significant correlation with pseudotime in both datasets (Hypergeometric test *p*-value < 1E-100), suggesting a core module of genes controls the indeterminate-determinate cell transition across ontogeny (Fig. 5B). In addition, pseudotime expression curves for identical genes in both datasets have a lower mean Fréchet distance than non-identical genes, indicating similar expression behavior across ontogeny (Fig. 5C). Among the 1003 shared genes, GO functions related to translation, cell wall, organ polarity, auxin, and gibberellin-related processes were enriched, likely reflecting the roles of auxin and gibberellic acid hormones in promoting differentiation in opposition to *KN1* that imposes indeterminacy (*2*, *25*). Furthermore, *BARELY ANY MERISTEM2*-*LIKE* (*BAM2-LIKE*), a *CLV1* paralog that regulates meristem and leaf development in Arabidopsis, shows upregulated expression among a subset of indeterminate cells as well as specific determinate cell populations, suggesting context-dependent signaling functions during maize shoot ontogeny (Fig. S9D-F) (*31*).

**Fig. 5.**
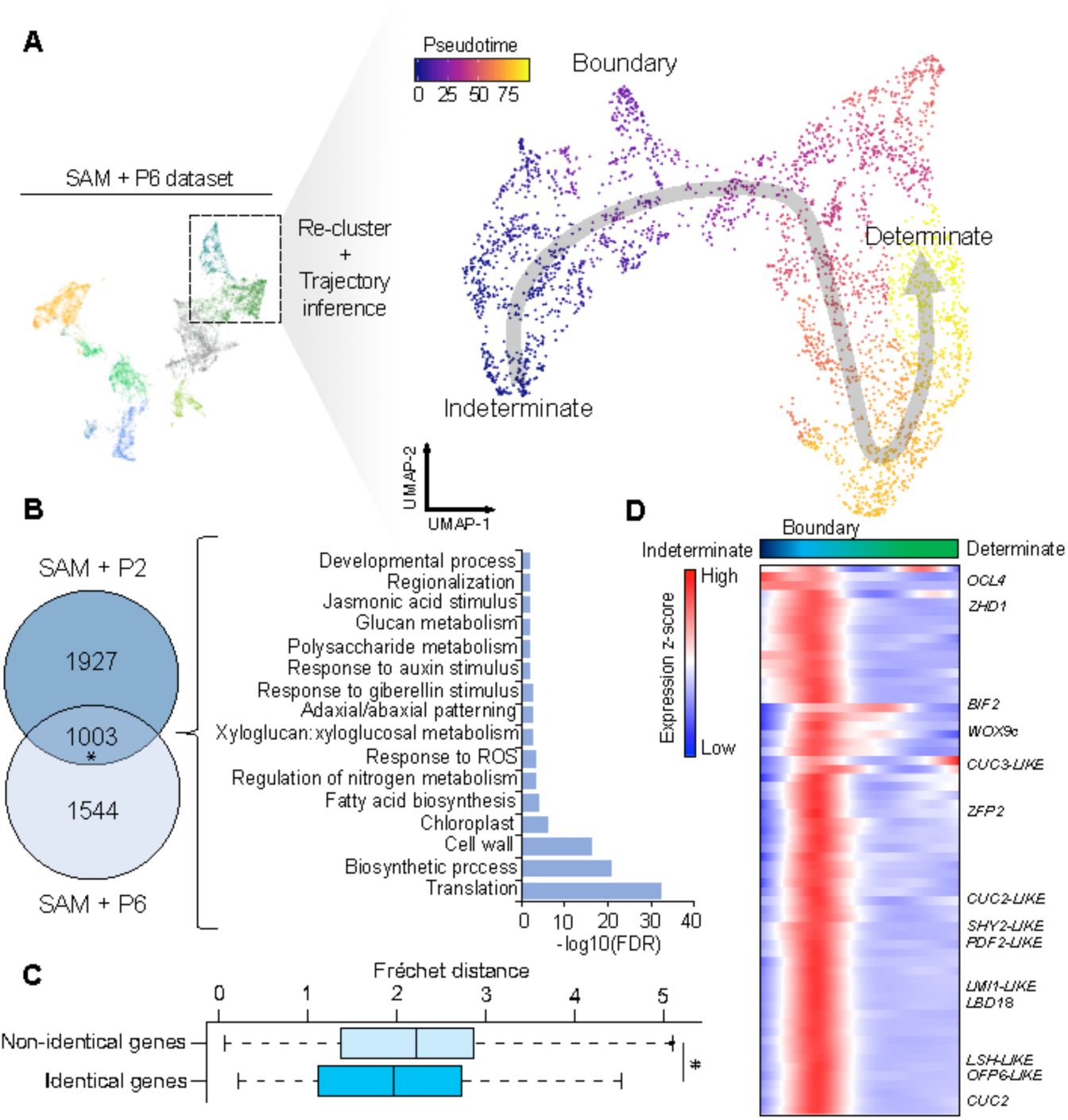
Identifying general features of the indeterminate-determinate cell fate transition across ontogeny. (A) A subset of cells from the SAM + P6 dataset (see Fig. S3B for inferred cluster identities) were re-clustered and trajectory inference was used to assign cell pseudotime scores along a transition from indeterminate to determinate cell fates. (B) Overlap of trajectory-correlated genes from the SAM + P2 and SAM + P6 datasets and their GO term enrichment (*, Hypergeometric test, *p*-value = 1.2E-320). (C) Average Fréchet distance for identical and non-identical genes in the SAM + P2 and SAM + P6 datasets (*, Wilcoxon rank-sum test, *p* = 5.2E-9). (D) Heatmap of genes with high specificity for a boundary region that delimits more indeterminate from more determinate cell fates with cells ordered by their pseudotime values.

Strikingly, the pseudotime trajectory analysis revealed a cell subpopulation at the transition between indeterminate and determinate cell fates, consistent with the persistence of a transcriptionally distinct developmental boundary (Fig S9A). Strong signatures of this boundary are evident in a subcluster marked by upregulated expression of several *CUC2*-like genes, which are known to promote organ separation in plants (Figure 5C) (*32*). Other genes specifically expressed in the subcluster include a putative *SHORT HYPOCOTYL2-LIKE* (*SHY2-LIKE*) gene that is boundary specific, which may balance auxin and cytokinin signaling to maintain undifferentiated fates (*33*). Likewise, *WOX9c* and a *LATE MERISTEM IDENTITY1-LIKE* (*LMI1-LIKE*) gene were also highly specific for boundary cells, along with a *LATERAL ORGAN BOUNDARIES DOMAIN-LIKE* (*LBD*-*LIKE*) and *LIGHT SENSITIVE HYPOCOTYLS-LIKE*(*LSH-LIKE*) genes. Taken together, the patterning of indeterminate to determinate cell identities in the shoot employs a shared set of gene functions during early and late phases of leaf ontogeny. Furthermore, the persistent boundary regions identified here transcriptionally partition indeterminate from determinate cell identities across ontogeny.

## Discussion

Here we present the first, to our knowledge, single-cell transcriptomic survey of a plant vegetative shoot apex including the stem cells housed within the SAM. This technique provides the key advantage of uncovering cell differentiation states and transitions in an unbiased fashion, without strict reliance on histological or genetic markers. Critically, we identify known and novel markers of the putative SAM stem-cell niche and show that it is characterized by increased expression of DNA methylation and DNA repair-related genes, as well as a low cell division rate. This observation supports the notion that only a subset of cells at the SAM tip have the specialized properties of stem cells, and may underlie the ability of plants to maintain high intergenerational genetic fidelity despite the absence of embryonic segregation of the germline as is found in animals (*8*). In addition, a low cell division rate and the upregulated expression of genes with genome protective functions highlights a shared convergently evolved solution to maintaining stem cells in plant and animals (*22*, *34*). Questions are raised regarding the lineage contributions of these stem cells to the developing plant. For example, do cells comprising the SAM tip make significant contributions to lateral organs, or are they preferentially destined to contribute to germ fates as described in the “*meristem d’attente*” (“*meristem in waiting*”) model proposed by early histological studies (*1*)? In the latter view, plant lateral organs are mostly derived from the transit-amplifying cell population that subtends the stem-cell domain, and the SAM tip makes minimal contributions to organogenesis. Emerging cell lineage tracing methodologies may shed light on these questions (*35*). Meanwhile, we find a lack of evidence for a WUS-expressing organizing center in the maize vegetative SAM, unlike what is modeled in other angiosperms, suggesting that genes with other, non-canonical, stem-cell organizing functions may be at play. Finally, by leveraging the continuum of cell states ranging from indeterminate to determinate cell identities, we reconstruct differentiation trajectories for cells of the seedling vegetative shoot apex. Many of the genes that are dynamically expressed along these pseudotime trajectories show similar accumulation patterns at early and later stages of leaf ontogeny, indicating the iterative redeployment of developmental programs.

## Methods

### Plant materials and growth conditions

Plants for single-cell RNA-Seq (scRNA-Seq), *in situ* hybridization, and phenotyping were grown in 72-well trays in a Percival A100 growth chamber under 16 hr days, a day temperature of 29.4 C, a night temperature of 23.9 C, and a relative humidity of 50%. Soil consisted of a 1:1 mixture of Turface MVP and LM111. The maize inbred B73 was used for scRNA-Seq and *in situ* hybridization analyses. The *gras32* and *gras33* alleles were obtained from the Maize Genetics Co-op Center (Urbana, IL, USA) in the W22 inbred background. Crosses were performed at a field site in Aurora, NY.

### In situ hybridization

RNA was isolated by Trizol extraction from liquid nitrogen-ground 2-week-old maize seedling shoot apices. Total RNA was DNAse I treated and cDNA was prepared using polyT-primed SuperScript III reverse transcriptase (Thermo Fisher Scientific, MA, USA) according to the manufacturer’s instructions. cDNA was then used as a template to amplify probe sequences, which were TA-cloned into the pCR4-TOPO vector backbone encoding flanking T3 and T7 polymerase promoters (Thermo Fisher Scientific) (See Table S10 for PCR primers). Following the verification of probe sequence and orientation by Sanger sequencing, antisense RNA probes were generated using a DIG-labeling kit and the resulting probes hydrolyzed as previously described (Roche Diagnostics, IN, USA) (*36*). An LNA probe was used for the *WOX9c in situ* presented in Figure 1H and was ordered directly from Qiagen (Hilden, Germany). LNA probe hybridization was carried out using 10 µM probe concentration and a 55 ° C hybridization temperature according to published methods (*37*).

Tissues for *in situ* hybridization were prepared and processed as previously described (*36*). Briefly, 2-week-old maize shoot apices were fixed overnight at 4°C in FAA solution (3.7% formalin, 5% acetic acid and 50% ethanol in water) and dehydrated through an ethanol series, cleared through increasing concentrations of Histo-Clear II (National Diagnostics, GA, USA), and then embedded in paraplast. 10 µm Sections were prepared using a Leica RM2235 microtome (Leica Biosystems, Wetzlar, Germany) and adhered to Probe-on-Plus microscope slides (Thermo Fisher Scientific). Sectioned tissues were then deparaffinized and rehydrated through a reverse ethanol series prior to Proteinase K treatment, refixation, and acetic anhydride treatment. Dehydrated tissues were then hybridized with probe overnight and washed, RNAse A treated, and incubated with AP-conjugated anti-DIG antibody (Roche Diagnostics). A colorimetric NBT/BCIP (Roche Diagnostics) reaction was then allowed to proceed until sufficient signal developed at which point the reaction was stopped in TE, slides were dehydrated, washed in Histo-Clear II, and mounted with Permount (Thermo Fisher Scientific). Images were obtained using an Axio Imager.Z10 (Carl Zeiss Microscopy, LLC, Thornwood, NY) microscope equipped with anAxioCam MRc5 camera.

### gras32 gras33 mutant analysis

Genomic DNA was extracted from the leaf tissue of F2 seedlings segregating for exonic gras32 and gras33 Mu insertion alleles. DNA extraction was performed as previously described (*38*). Plants were genotyped by PCR (see Table S10 for primers) and Paraffin-embedded FAA-fixed shoot apex tissues (see *in situ* hybridization) were longitudinally sectioned to 10 µm and adhered to Probe-on-Plus slides. Tissues were deparaffinized and rehydrated through an ethanol series and then equilibrated in 1% sodium borate (w/v). Tissues were then stained in a 0.5% solution of *o*-Toluidine (TBO) in 1% sodium borate for 5 min followed by an ethanol dehydration series, washing in Histo-Clear II, and mounting in Permount. Samples were then imaged (see *in situ* hybridization). SAM width and height were determined in medial sections using ImageJ.

### Cell cycle quantification

Medial SAM sections probed by *in situ* hybridization for expression of the S-phase and G2/M-phase upregulated H3 and CYC1 genes, respectively, were imaged and imported into ImageJ. Images were converted to 16-bit format and processed using the thresholding tool such that stained areas could be differentiated from non-stained areas. For each SAM, five sections of equal height were measured and the percent ratio of above threshold (stained) area relative to the area of each bin was calculated.

### Generation and collection of protoplasts

Two-week-old maize seedlings were hand dissected to either a ~3 mm portion of the stem and plastochron 6 primordia (SAM + P6) or the SAM and plastochron 2 (SAM + P2). Dissected tissue was briefly macerated and placed immediately in protoplasting solution, which consisted of 0.65 M mannitol, 1.5% Cellulase R10, 1.5% Cellulase RS, 1.0% Macerozyme, 1.0% hemicellulase (Sigma-Aldrich), 10 mM MOPS pH 7.5, 10 mM L-Arginine HCl pH 7.5, 1 mM CaCl2, 5 mM ß-mercaptoethanol, and 0.1% BSA. Prior to the addition of CaCl2, ß-mercaptoethanol, and BSA, the solution was heated to 55°C for 10 min to facilitate enzyme solubilization. All tissue collection was completed within 30-45 min. Tissue digestion was carried out with gentle shaking at 29°C for 2 hrs. After digestion, fluorescein diacetate (FDA) was added to the cell suspension at a concentration of 5 μg/mL and cells were allowed to incubate in darkness for 5 min. The cell suspension was then filtered using a 40 μm nylon filter and the cells were centrifuged at 250 × g for 3 min at 4°C. Cells were resuspended in washing buffer consisting of 0.65 M mannitol, 10 mM MOPS pH 7.5, and 10 mM L-Arginine pH 7.5 and washed 3 times using the same centrifugation conditions. Cell viability and concentration were assessed using a hemocytometer and a fluorescent Axio Imager.Z10 microscope equipped with a 488 nm filter for FDA staining detection.

Cells isolated from SAM + P6 tissue were suspended at a concentration of ~10,000 cells/mL and loaded onto the 10X Genomics Chromium Controller using v3 reagents following the manufacturer’s instructions to target approximately 10,000 cells. Cells from the SAM + P2 tissue were resuspended in 1 mL of wash buffer and a 100-200 uL aliquot was transferred to the well of a clear-bottom CoStar plate kept over ice. 200 uL of wash buffer was distributed to other wells on the plate. A Leica M205 FCA microscope equipped with a 488 nm fluorescent filter was then used to transfer individual viable (FDA+) cells. Each cell was carried in 0.1 μL volumes through three wash buffer wells. Following washing, cells were transferred to a 96-well LoBind (Eppendorf) plate containing reagents for reverse transcription (see library construction and sequencing), kept on dry ice. Cell collection was completed in <1 hr. Plates were sealed with adhesive film and transferred to a −80°C freezer prior to further processing.

### Cell viability assays

Protoplasts were obtained as described (see generation and collection of protoplasts) with the exception of the pH and concentration of buffer components used. For the pH 5.7 condition, 10 mM MES buffer was used whereas for the pH 6.5, 7.0, and 7.5. conditions 10 mM MOPS buffer was used. To quantify cell viability, the ratio of fluorescing (living) protoplasts to dead (non-fluorescing) and large debris (> 5 μm) were quantified in each of the four larger corners of a hemocytometer using a fluorescent Axio Imager.Z10 microscope equipped with a 488 nm filter.

### SAM + P2 single-cell RNA isolation and amplification

Single-cell RNA-Seq library construction was performed using an adaptation of the Cel-Seq2 protocol (*39*). 96-well LoBind plates were prepared with each well containing 0.22 μL 25 ng/μL Cel-Seq2 RT primer (1s – 96s, Supplementary Table 10), 0.11 μL 10 mM dNTPs, and 0.77 μL nuclease-free H_2_O such that each well of the plate contained a unique cell barcoded RT primer. After cell collection and storage at −80°C, plates were thawed briefly on ice and centrifuged at 2000 × *g* for 2 min at 4°C. Next, plates were incubated at 65°C for 5 min and again centrifuged using the same settings. For reverse transcription, 0.54 μL First Strand Buffer, 0.27 μL 0.1 M DTT, 0.135 μL RNAseOUT (Thermo Fisher Scientific), 0.034 μL SuperScript III, and 0.52 μL nuclease-free H_2_O were added to each well and plates were incubated for 1 hr at 42°C followed by RT inactivation for 10 min at 70°C. cDNA was pooled horizontally into the eight wells at the end of each plate. Then, 2.5 μL 10X Exonuclease I buffer and 2.1 μL Exonuclease I were added to each well and incubated at 37°C for 15 min followed by heat inactivation at 80°C for 15 min. cDNA was purified using Ampure Clean XP beads according to the manufacturer’s instructions and resuspended in 7 μL nuclease-free H_2_O. Second strand synthesis was performed by adding 2.31 μL Second Strand Buffer, 0.23 μL dNTPs, 0.08 μL *E. coli* DNA Ligase, 0.3 μL *E. coli* DNA Polymerase, and 0.08 μL RNAse H and incubated at 16°C for 2 hrs. Pooled dsDNA from each of the eight reactions was pooled and then purified using Ampure XP beads according to the manufacturer’s instructions followed by resuspension in 6.4 μL nuclease-free H_2_O. *In vitro* transcription was performed by the addition of 1.6 μL each of the A, G, C, U dNTPs, 10X T7 Polymerase Buffer, and T7 Polymerase (Ambion, TX, USA) followed by incubation at 37°C for 13 hrs. Amplified RNA was then treated with ExoSAP-IT (Thermo Fisher Scientific) for 15 min at 37°C followed by RNA fragmentation by addition of 5.5 μL fragmentation buffer (200 mM Tris pH 8.0, 150 mM MgCl_2_) and incubation for 3 min at 94°C. Fragmentation was stopped by transfer to ice and the immediate addition of 2.75 μL 0.5 M EDTA pH 8.0.

### GLibrary construction and sequencing

For reverse transcription, 5 μL of the amplified RNA was added to 1 μL randomhexRT primer and 0.5 μL dNTPs, incubated at 65°C for 5 min, and chilled on ice. Next, 2 μL First Strand Buffer, 1 μL 0.1 M DTT, 0.5 μL RNAseOUT, and 0.5 μL SuperScript III were added and the samples incubated for 10 min at 25°C followed by a 1 hr incubation at 42 °C. Half of the completed RT reaction was subjected to PCR by addition of 5.5 μL nuclease-free H_2_O, 12.5 μL Phusion High-Fidelity PCR Master Mix with HF Buffer (New England Biolabs, MA, USA), RNA PCR Primer1 (RP1), and 1 μL of RNA PCR Primer X (RPIX). PCR conditions were as follows: 30 s at 98°C, 11 cycles of: [10 s at 98°C, 30 s at 60°C, 30 s at 72°C], and 10 min at 72°C. The pooled cDNA from each plate received a unique RPIX. Samples were then purified and size-selected via two rounds of AMPure XP bead treatment using a 1:1 ratio of beads-to-sample and the final library resuspended in 10 μL nuclease-free H_2_O. 1 μL of the library was submitted for fragment analysis using a Bioanalyzer to confirm a target library size between 200-400 bp. An additional 1 uL was used for concentration measurement using a Qubit. If the library concentration was suboptimal, the second unused half of the RT reaction was amplified using up to 15 PCR cycles. Libraries were then sequenced using a single lane on an Illumina NextSeq 500 instrument using the small RNA chemistry. Paired-end sequencing was performed with 15 and 77 bp obtained for read 1 and read 2, respectively. The libraries generated from the biological replicates of SAM + P6 cells were also sequenced using a NextSeq 500 instrument, with each replicate allocated a single lane of sequencing.

### Single-cell RNA-Seq read processing and cell filtering

SAM + P2 FASTQ files were processed using the default settings in the celseq2 pipeline (https://github.com/yanailab/celseq2), which includes read trimming, alignment, and UMI counting steps to generate a UMI count matrix (*39*). Reads were aligned to version 3 of the B73 reference genome. SAM + P6 reads were trimmed, aligned, and UMI count matrices generated using the CellRanger version 3.1.0 pipeline under the default settings. Reads were aligned to version 3 of the B73 reference genome. The UMI count matrices for individual biological replicates were merged prior to further analysis. For the SAM + P2 dataset, cells with fewer than 500 genes detected were removed while in the SAM + P6 dataset, cells with fewer than 2500 genes detected were removed. In both datasets, cells with over 1% of transcripts encoded by the mitochondrial genome were removed.

### Dimensionality reduction, cell type classification, and differential expression analysis

Cell type analysis and clustering were performed using Seurat v3.0 (*40*). The merged UMI count matrices were converted to Seurat objects. Normalization and variance stabilization were performed using SCTransform and the 3000 genes with the highest expression variability were used for the calculation of principal components. Uniform Manifold Approximation and Projection (UMAP) was then used to embed cells in lower dimensional space for data visualization. For projection of SAM + P2 cells, UMAP was run using dim = 1:5, n.neighbors = 15, min.dist = 0.1, and spread = 5. For the projection of SAM + P6 cells, UMAP was run using dim = 1:25, n.neighbors = 25, min.dist = 0.01, and spread = 1. For the SAM+P6 subset cells, cells belonging to clusters 5 and 0 were re-clustered in isolation using the same parameters as for the full dataset. Cells were assigned to clusters using k-means hierarchical clustering. All differential expression analyses used to compare gene expression on a per-cluster basis were performed using Wilcoxon ranked-sum tests. Gene ontology (GO) enrichment analysis was done using a Fisher’s exact test implemented in AgriGO v2 (*41*). Cell cycle regression was used to reduce the effects of the cell cycle on cell clustering in the SAM + P2 dataset. Differentially expressed genes among cells belonging to cell clusters with S-phase and G2/M-phase marker gene expression were first identified (adjusted p-value < 0.05). Genes that were highly specific for these clusters were identified using a ratio of the number of cells expressing a given differentially expressed gene within the cluster to those in all clusters. Those with a ratio greater than 2 were deemed phase specific and their expression was used to calculate a numeric cell cycle score and a cell cycle stage (G1, S, G2/M). SCTransform was run again on the raw UMI count matrix with cell cycle score as a variable to regress, followed by PCA and UMAP dimensionality reduction.

### Trajectory inference and pseudotime analysis

Trajectory inference and pseudotime analysis was performed using Monocle3 (https://cole-trapnell-lab.github.io/monocle3) (*42*). A principal graph was generated using the learn_graph function and cells were assigned a pseudotime value using order_cells with a pseudotime start or “root” position manually selected. Genes that were differentially expressed along the inferred trajectories were identified using the graph_test function, which applies a Moran’s *I* test to detect spatial autocorrelation. For the SAM+P2 dataset, individual Moran’s *I* tests were performed on cell subsets to better identify genes with branch-specific expression patterns. This involved two tests on a common population of cells derived from the Tip, Meristem 1, and Meristem 2 clusters merged with cells from the Primordia and Vasculature clusters or the Epidermis 1 and 2 clusters. The significantly differentially expressed genes along both trajectories were then further analyzed. For the visualization and analysis of pseudotime-dependent gene expression patterns, cubic smoothing splines were fit to each gene using the R smooth.spline function with a spar parameter of 1.1. To compare the expression behavior of genes in the SAM+P2 and SAM+P6 datasets, smoothed expression profiles for each gene were averaged in 10 pseudotime bins and z-z-scaled values for each bin were calculated. The Fréchet distances between the curves of identical genes and all non-identical genes were calculated using the SimilarityMeasures R package (*43*).

### Phylogenetic analysis

For the phylogenetic analysis of GRAS proteins, amino acid sequences of all Arabidopsis GRAS proteins and HAM-LIKE homologs in rice (*Oryza sativa*) and maize were downloaded from Phytozome. Amino acid sequences were then aligned using Clustal Omega. Maximum likelihood phylogenetic tree construction was performed using PhyML. The Jones-Taylor-Thornton (JTT) amino acid substitution model was selected based on its Akaike Information Criterion (AIC) calculated using Smart Model Selection (SMS) implemented in PhyML (*44*). Branch support values were calculated using the aLRT SH-LIKE fast likelihood-based method (*45*).

## Acknowledgments

We thank the Cornell University Biological Resource Center for assistance in single-cell genomics and sequencing techniques as well as J. Cammarata and A. Roeder for critical feedback on the manuscript. This work was funded by National Science Foundation grants IOS-2016021 and IOS-1238142 (to M.J.S) and a Schmittau-Novak Small Grant (to J.W.S.).

## Author contributions

J.W.S. and M.J.S. conceptualized the study, J.W.S analyzed resulting data, J.W.S. and J.S. performed experiments, J.W.S. and M.J.S wrote the manuscript with J.S. contributing to review and editing.

## Data Availability

Sequencing data generated in this work are available at the NCBI Short Reads Archive (accession no. PRJNA637882).

## Additional Information / Supplementary Materials

Supplementary Figures S1-S9

Supplementary Datasets S1-S10

**Fig. S1.**
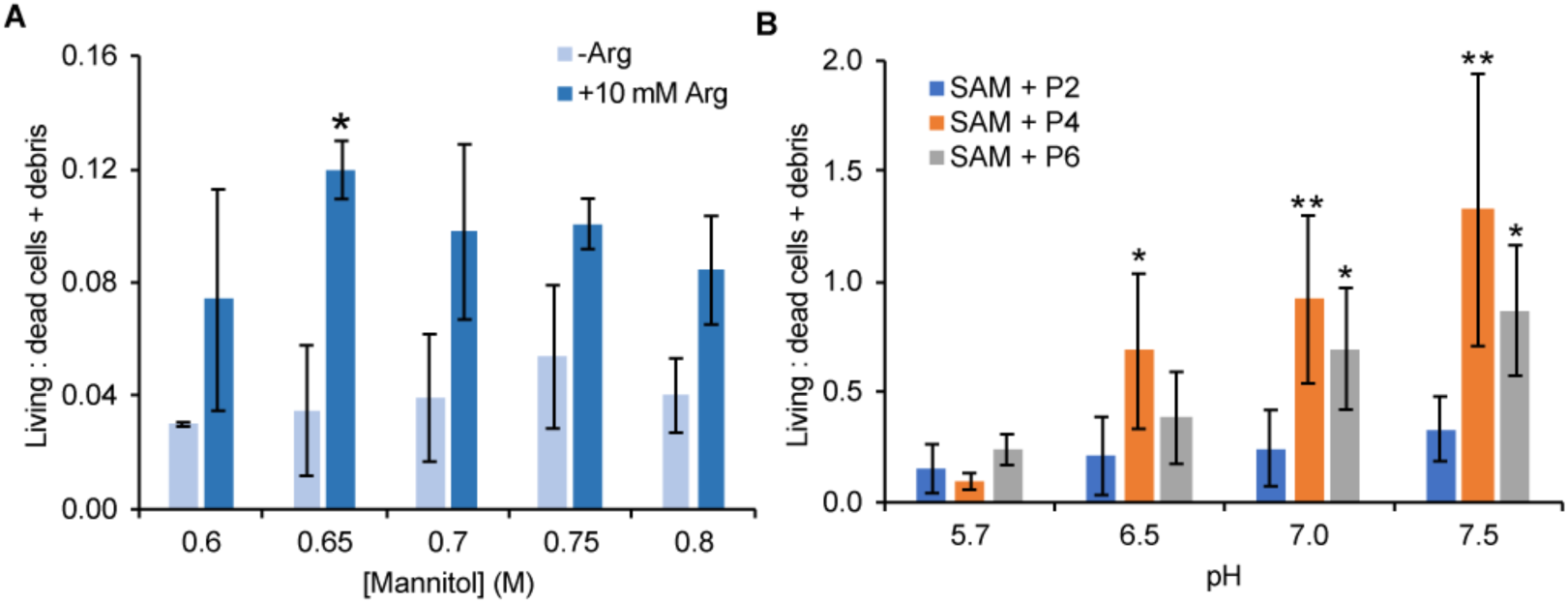
L-Arginine supplementation and increased pH improve maize protoplast viability. (A) Ratio of FDA+ (viable) protoplasts relative to dead cells and large (> 5 µm) cellular debris in protoplasts isolated from SAM+P4 tissue carried out in solutions with varying mannitol and 10 mM L-arginine supplementation (n = 2) Asterisks signify the result of a Student’s t-test comparing the mean of -Arg and +10 mM Arg samples at each pH value (*, *p* < 0.05). (B) Protoplast viability assayed in solutions buffered at varying pH levels and supplemented with 10 mM L-arginine from three different tissue types (n = 2-3). Asterisks signify the result of a Student’s t-test comparing the mean of each tissue type at a given pH compared to its mean at pH 5.7 (*, *p* < 0.05; **, p < 0.01).

**Fig. S2.**
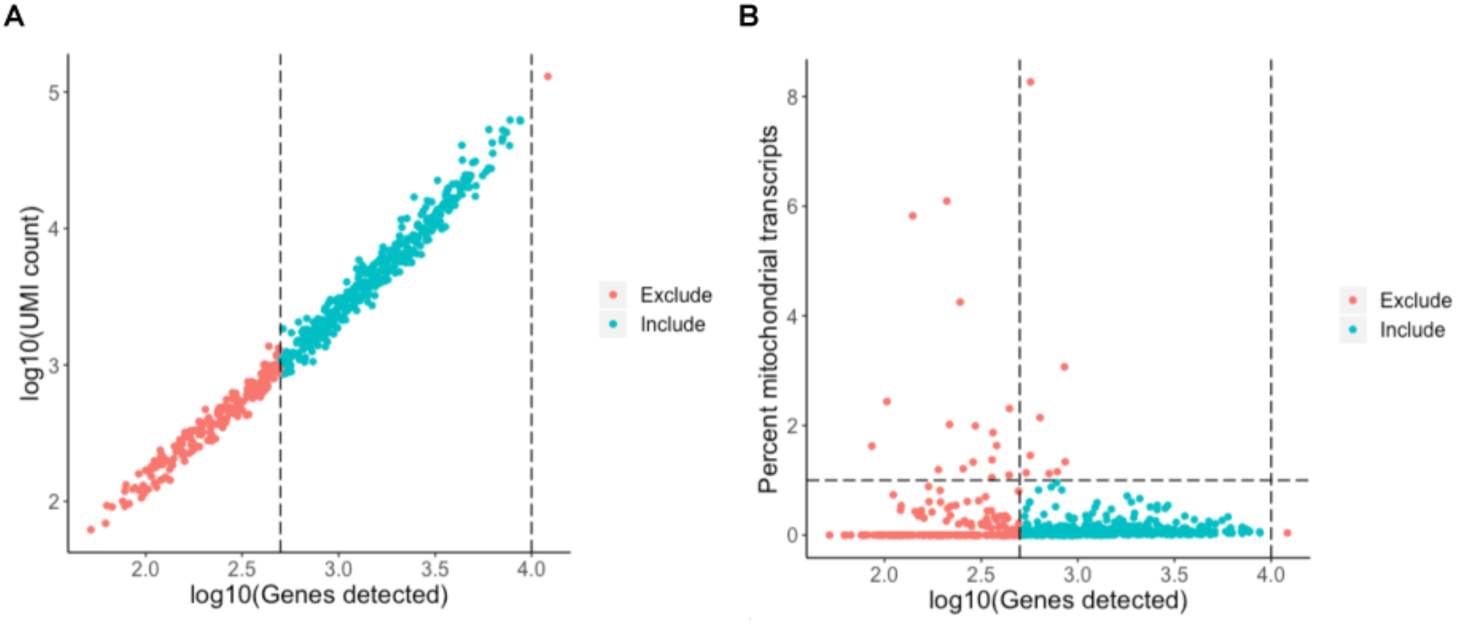
Cell filtering for the SAM+P2 dataset. (A) Relationship between the number of genes and transcripts detected per cell in the SAM+P2 dataset. The dashed line indicates the genes per cell cutoff used to filter low quality cells. (B) Relationship between the number of genes detected and the percentage of mitochondrial transcripts per cell in the SAM+P2 dataset. Dashed lines indicate the genes per cell and percent mitochondrial transcripts filtering cutoffs.

**Fig. S3.**
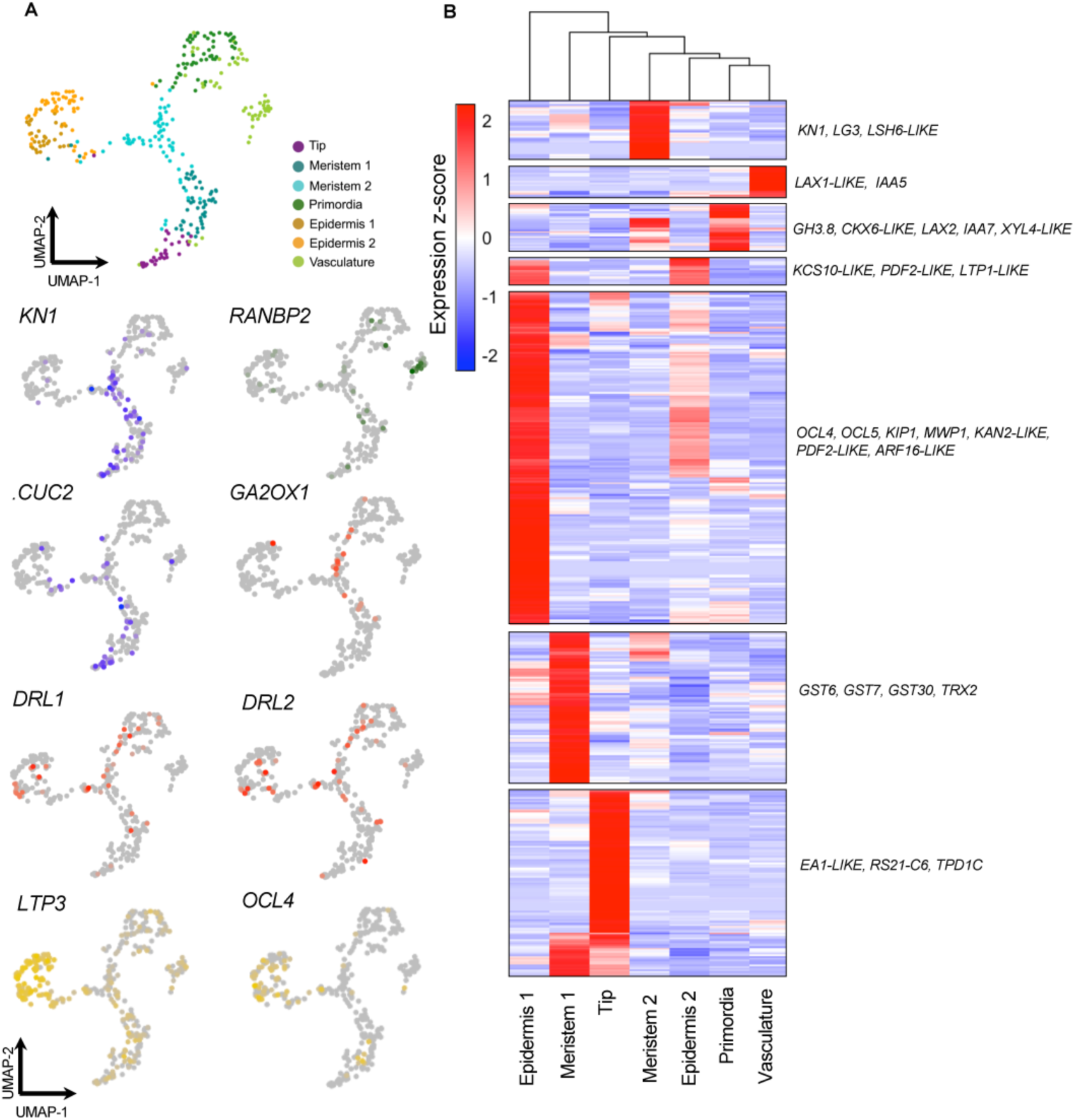
Expression patterns of markers used for cell type inference. (A) Dimensionality reduction of the SAM + P2 dataset and associated marker genes used to infer cell identity. (B) Heatmap of all identified marker genes for each cell type cluster with representative markers of each group indicated to the right.

**Fig. S4.**
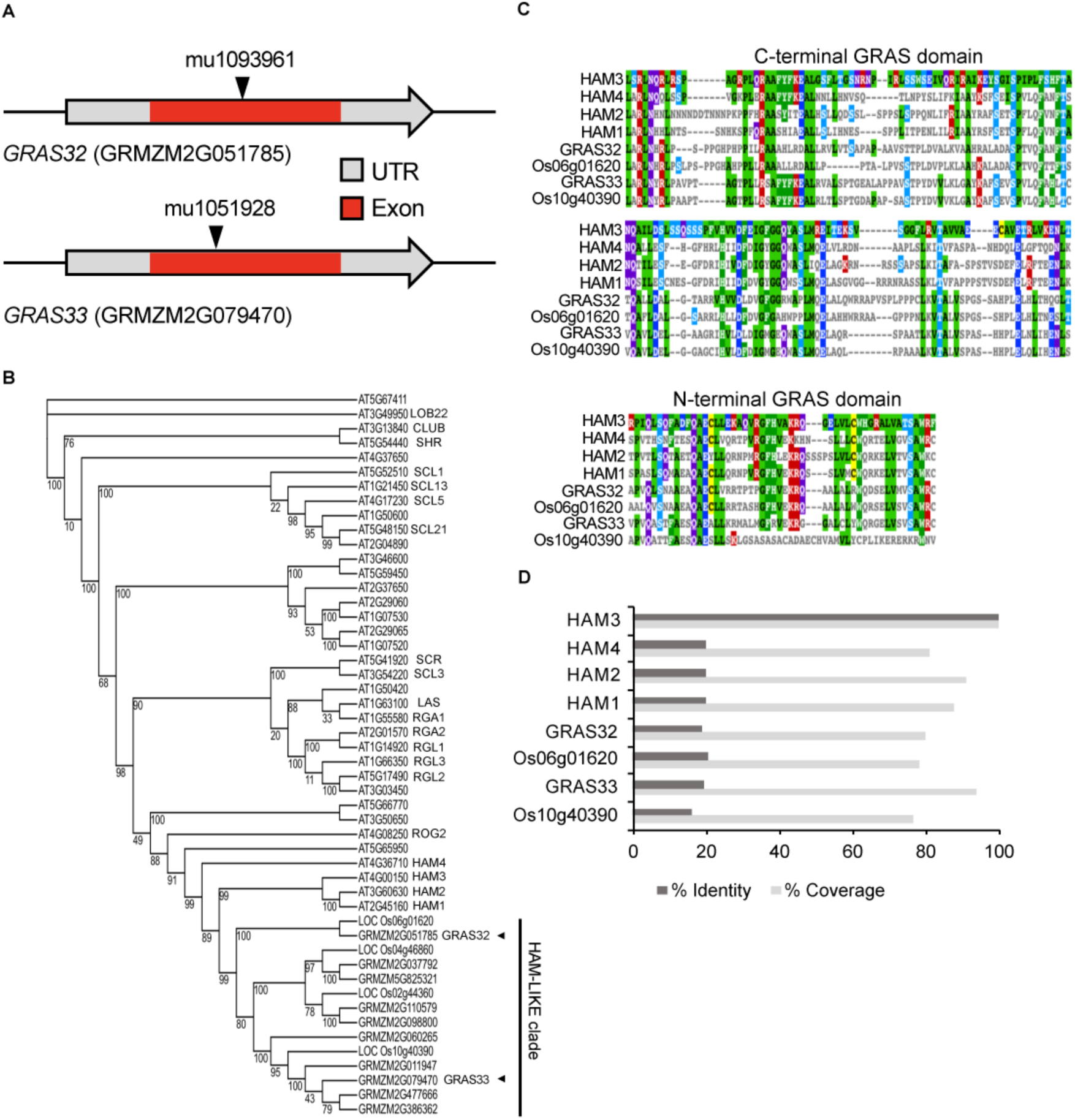
Maize *GRAS32* and *GRAS33* are *HAM-LIKE* genes. (A) Gene models for maize *GRAS32* and *GRAS33* showing the *Mutator* transposon insertion sites for the mutant alleles used in this study. (B) Maximum likelihood phylogenetic tree of Arabidopsis GRAS family transcription factor amino acid sequences with maize and rice HAM-LIKE homologs. Branch support values are the result of the Approximate Likelihood-Ratio test. (C) Amino acid alignment showing regions of high similarity in the N- and C-terminal regions of the GRAS domain between select maize, rice, and Arabidopsis HAM proteins. (D) Amino acid percent identity and percent coverage of HAM homologs relative to Arabidopsis HAM3.

**Fig. S5.**
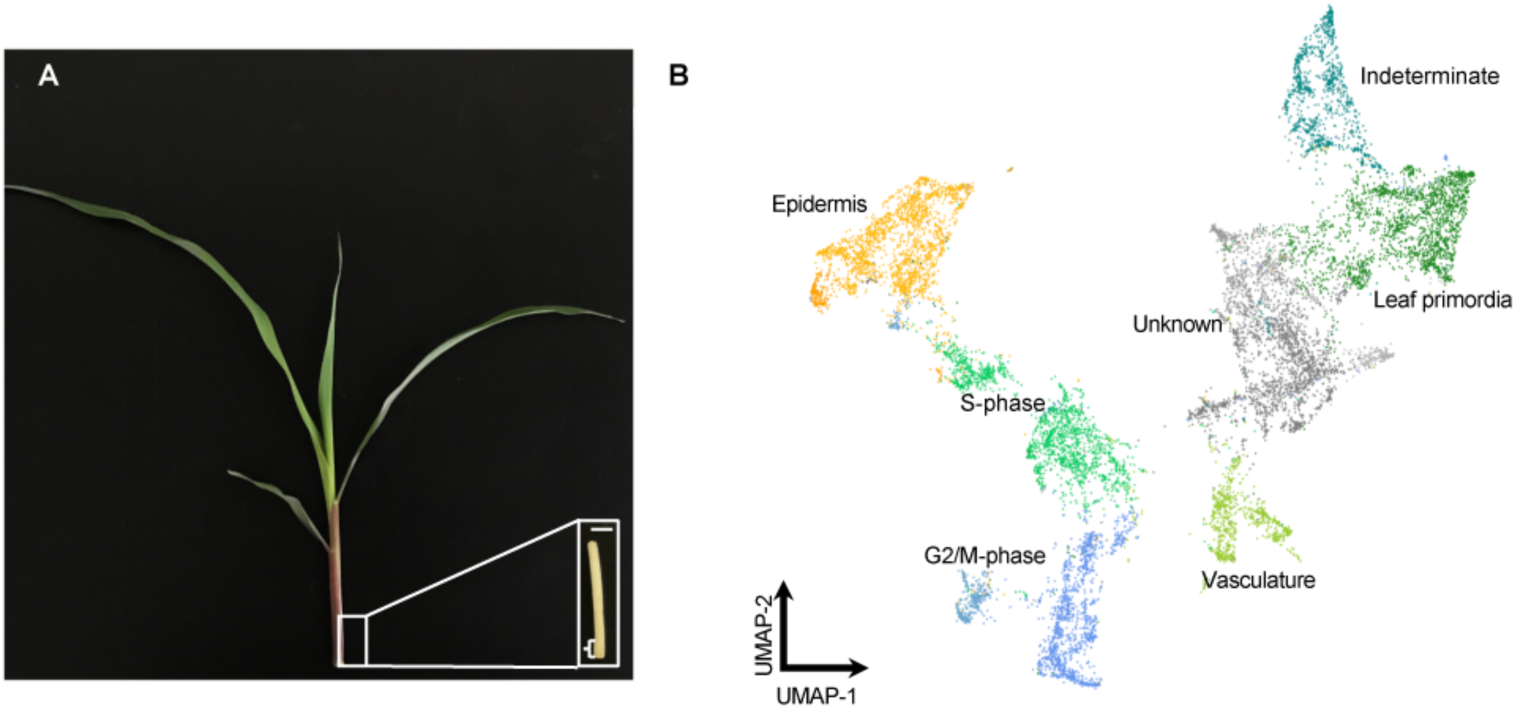
Tissue collection and cell type representation in the SAM+P6 dataset. (A) Two-week-old inbred B73 maize seedling. (Inset) Dissection to plastochron 6 (P6) showing the tissue targeted for analysis consisting of approximately 3 mm of stem and P6 tissue (scale bar = 5 mm). (B) Dimensionality reduction of the filtered cellular transcriptomes represented in the SAM + P6 dataset along with their inferred cluster identities.

**Fig. S6.**
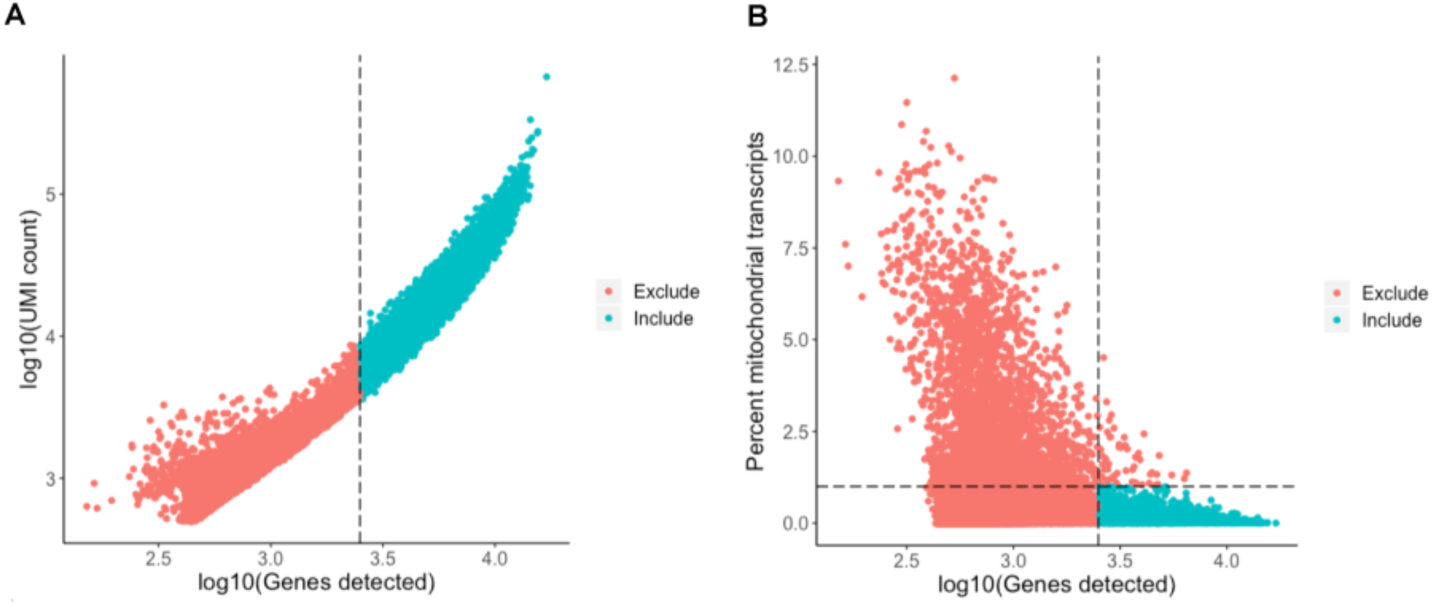
Cell filtering for the SAM+P6 dataset. (A) Relationship between the number of genes and transcripts detected per cell in the SAM+P6 dataset. The dashed line indicates the genes per cell cutoff used to filter low quality cells. (B) Relationship between the number of genes detected and the percentage of mitochondrial transcripts per cell in the SAM+P6 dataset. Dashed lines indicate the genes per cell and percent mitochondrial transcripts filtering cutoffs.

**Fig. S7.**
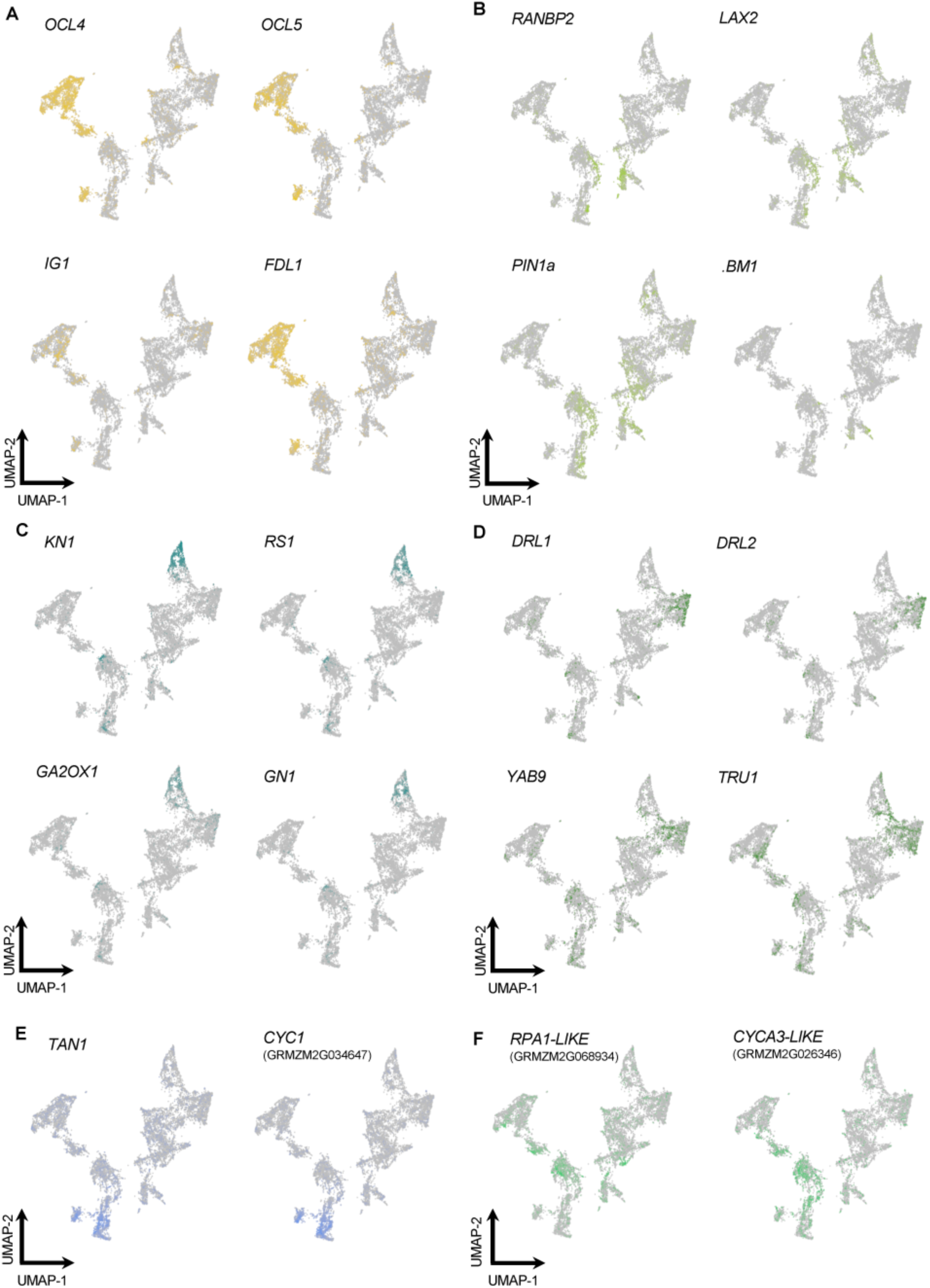
Expression patterns of markers used for cell type inference. Selected marker genes for: (A) epidermis;. (B) vasculature; (C) indeterminate (stem tissue); (D) leaf primordia; (E) G2/M-phase; (F) S-phase.

**Fig. S8.**
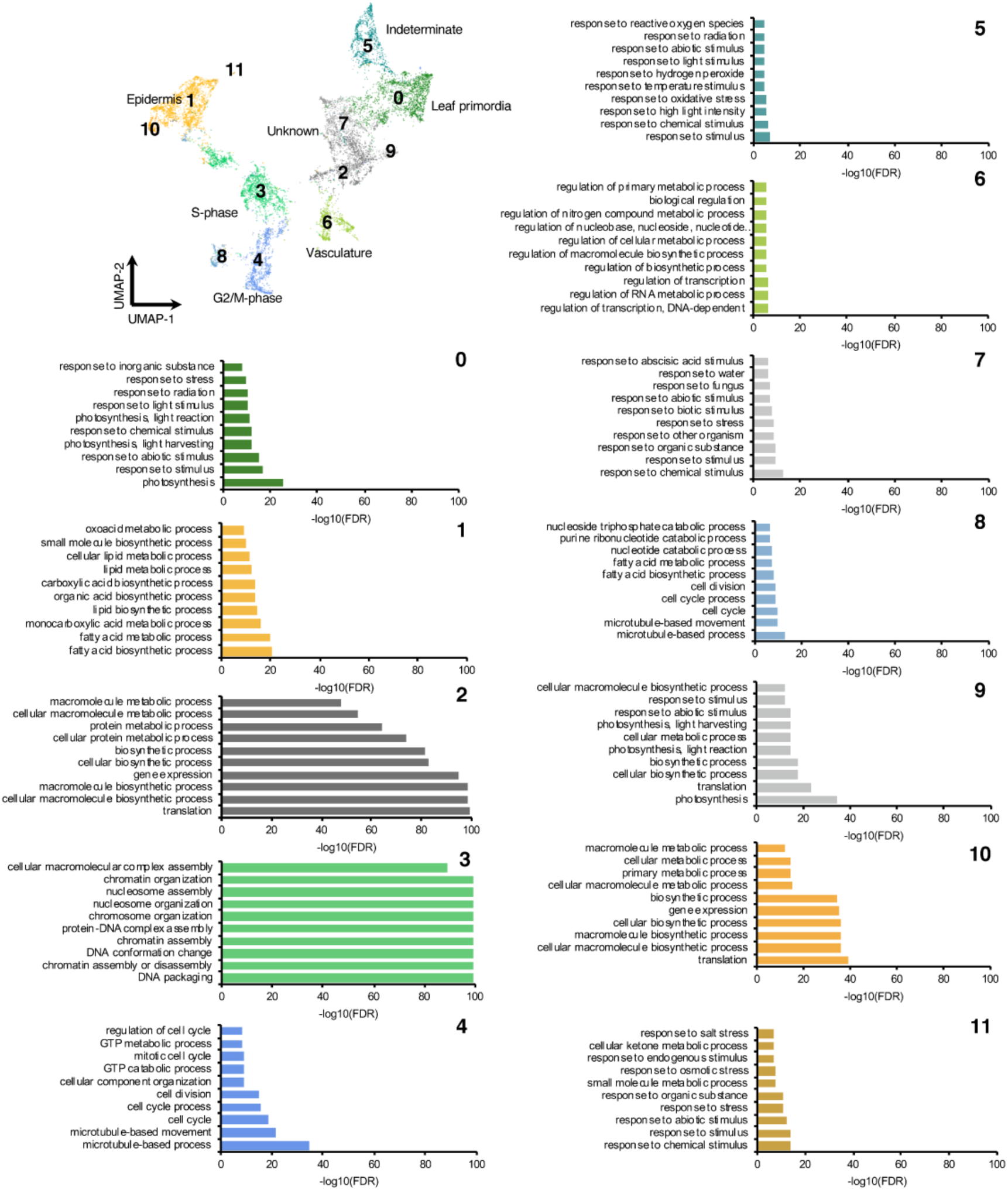
Gene ontology (GO) enrichment for each of the identified clusters in the SAM + P6 dataset. Clusters identified by hierarchical clustering were numbered. Bar plots display the top 10 most significant GO enrichments for genes that mark each cluster.

**Fig. S9.**
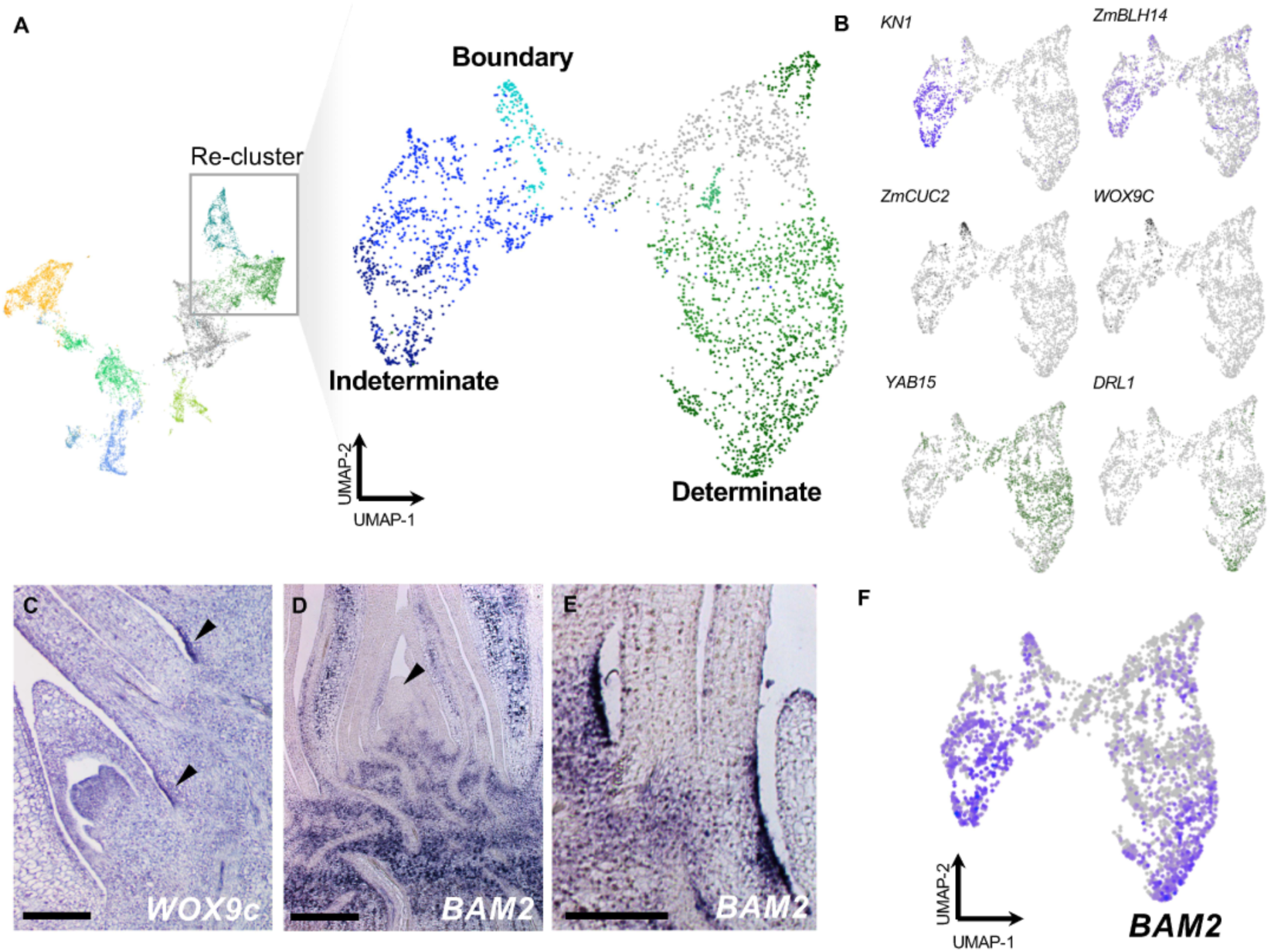
Marker gene expression among cells with determinate and indeterminate identity. (A) Re-clustering and cluster assignment of cells identified in the SAM + P6 dataset. (B) Expression patterns of marker genes among re-clustered cells. (C) *WOX9c* transcript accumulation patterns in boundary regions of the seedling shoot system (arrowheads). (D,E) *BAM2* expression patterns in the seedling shoot showing expression in indeterminate and determinate cell populations (with the SAM indicated by an arrowhead) (D) as well as in boundary regions (E). (F) *BAM2* expression in the SAM + P6 dataset. Scale bars = 250 *µ*m.

**Table S1. Genes used for cell cycle regression.**

**Table S2. Differentially expressed genes for all clusters in the SAM + P2 dataset.** Statistical results (*p*-value) reflect the output of a Wilcoxon ranked sum test followed by a Bonferonni correction (Adjusted *p-*value) performed in Seurat v3.

**Table S3. Gene Ontology (GO) term enrichment for differentially expressed genes in each SAM + P2 cluster.** Differentially expressed genes (see Table S1, p. adj < 0.05) were analyzed for GO Term enrichment using AgriGo v2.

**Table S4. Average cluster-based expression of potential maize CLV1-CLV3-WUS pathway genes.** Values reflect average gene expression in each cluster (UMI counts).

**Table S5. Differentially expressed genes in cells positive for expression of the core marker gene (GRMZM2G049151) in the SAM + P2 dataset.** Statistical results (*p*-value) reflect the output of a Wilcoxon ranked sum test followed by a Bonferonni correction (Adjusted *p-*value) performed in Seurat v3.

**Table S6. Genes with dynamic expression over pseudotime in the SAM + P2 dataset.** Statistical results reflect the output of a Moran's *I* test implemented in Monocle v3 for epidermal and primordia/vasculature trajectories.

**Table S7. Differentially expressed genes for all clusters in the SAM + P6 dataset.** Statistical results (*p*-value) reflect the output of a Wilcoxon ranked sum test followed by a Bonferonni correction (Adjusted *p-*value) performed in Seurat v3 (.

**Table S8. GO Term enrichment for heatmap clusters of pseudotime-correlated genes.** Differentially expressed genes (see Table S7, p. adj < 0.05) were analyzed for GO Term enrichment using AgriGo v2.

**Table S9. Gene Ontology (GO) term enrichment for differentially expressed genes in each SAM + P6 cluster.** Differentially expressed genes (see Supplementary Table 1, p. adj < 0.05) were analyzed for GO Term enrichment using AgriGo v2.

**Table S10. Oligonucleotide and primer sequences used in this study.**

